# Theory of measurement for site-specific evolutionary rates in amino-acid sequences

**DOI:** 10.1101/411025

**Authors:** Dariya K. Sydykova, Claus O. Wilke

## Abstract

In the field of molecular evolution, we commonly calculate site-specific evolutionary rates from alignments of amino-acid sequences. For example, catalytic residues in enzymes and interface regions in protein complexes can be inferred from observed relative rates. While numerous approaches exist to calculate amino-acid rates, it is not entirely clear what physical quantities the inferred rates represent and how these rates relate to the underlying fitness landscape of the evolving proteins. Further, amino-acid rates can be calculated in the context of different amino-acid exchangeability matrices, such as JTT, LG, or WAG, and again it is not well understood how the choice of the matrix influences the physical interpretation of the inferred rates. Here, we develop a theory of measurement for site-specific evolutionary rates, by analytically solving the maximum-likelihood equations for rate inference performed on sequences evolved under a mutation–selection model. We demonstrate that for realistic analysis settings the measurement process will recover the true expected rates of the mutation–selection model if rates are measured relative to a naïve exchangeability matrix, in which all exchangeabilities are equal to 1/19. We also show that rate measurements using other matrices are quantitatively close but in general not mathematically equivalent. Our results demonstrate that insights obtained from phylogenetic-tree inference do not necessarily apply to rate inference, and best practices for the former may be deleterious for the latter.

**Significance Statement:** Maximum likelihood inference is widely used to infer model parameters from sequence data in an evolutionary context. One major challenge in such inference procedures is the problem of having to identify the appropriate model used for inference. Model parameters usually are meaningful only to the extent that the model is appropriately specified and matches the process that generated the data. However, in practice, we don’t know what process generated the data, and most models in actual use are misspecified. To circumvent this problem, we show here that we can employ maximum likelihood inference to make defined and meaningful measurements on arbitrary processes. Our approach uses misspecification as a deliberate strategy, and this strategy results in robust and meaningful parameter inference.

---

**A** quantity of broad interest in molecular evolution is the site-specific rate of evolution, defined as the rate at which a given nucleotide, amino-acid, or codon position evolves in a gene or genome (1). Site specific evolutionary rates can be used, for example, to identify sites of structural or functional importance in proteins, as those sites tend to evolve at the lowest rates (2–10). Several methods have been proposed to measure site-wise rate of evolution in protein sequences (11–18). These methods all measure rates as scaling factors in front of a fixed exchangeability matrix describing the relative likelihood that certain amino acids are substituted by other ones. The exchangeability matrix is usually not estimated from the data, because it contains many more parameters than can be reliably estimated from moderately sized alignments. Significant effort has been expended into empirically deriving exchangeability matrices from large sequence databases (19–27). However, recent analysis has shown, somewhat surprisingly, that rate estimates are largely insensitive to the choice of exchangeability matrix used in the inference process (16). And yet, a simplistic equal-rates exchangeability matrix could uniquely identify rapidly-evolving sites that models with empirically-derived exchangeabilities could not (16). These observations highlight that the optimal choice of the exchangeability matrix is both non-obvious and consequential in practical applications.

The rate parameters of interest are generally estimated via maximum likelihood or Bayesian methods, and they are expressed as scalar variables inside the model likelihood function. While the estimation process for such scalar parameters is well understood from a statistical modeling perspective, the statistical perspective does not provide insight into the physical meaning of the estimated rate parameters. What actual physical quantities do the parameter estimates represent? To make progress on this question, it helps to think of the model fitting process as a measurement process. By fitting a likelihood model to an alignment of sequence data, we are choosing numbers (the parameter estimates) that are in some quantifiable way linked to the reality that created the data. Importantly, how changes in the real world are reflected in the estimated parameters is not obvious from inspecting the likelihood function. It requires a careful calculation of the expected parameter estimates given specific underlying realities. The link between a parameter estimate and the real world is called a theory of measurement (28), and we develop here such a theory for site-specific evolutionary rates in amino-acid sequences.

Our overall approach is to assume we know the process that generates the data (i.e., describes the real world), and then we use maximum likelihood to calculate evolutionary rates under different estimation procedures. We show that this procedure can be performed analytically for a broad class of generating models and it produces meaningful closed-form solutions. Our analysis reveals that by measuring rates with a naïve, equal-rates exchangeability matrix we can recover the true expected rates of the generating process. Thus, in a precise, quantifiable sense, we can state that the naïve exchangeability matrix is the ideal matrix for performing meaningful measurements of evolutionary rates. Our analysis also explains why commonly used exchangeability matrices yield rate inferences that are similar to each other and to the naïve substitution matrix.

## Results

The evolution of a genetic sequence can be thought of as a single ancestor sequence changing over time and diverging into multiple descendant sequences. Different sites evolve at different rates (1), which will cause the descendant sequences to vary at some sites from each other and from the ancestor sequence (Figure 1A). The evolutionary divergence of sequences over time is commonly modeled with a continuous-time Markov model (29), and most models currently in practical use assume that each site evolves independently and thus can be described by its own Markov chain (but see (30, 31)). A mutation may occur at any site, and it subsequently either fixes or is lost to drift. The Markov model simplifies the substitution process by capturing only the mutations that fix (32–35), and the states of the Markov chain correspond to the set of possible mutations that can arise and fix.

**Fig. 1.**
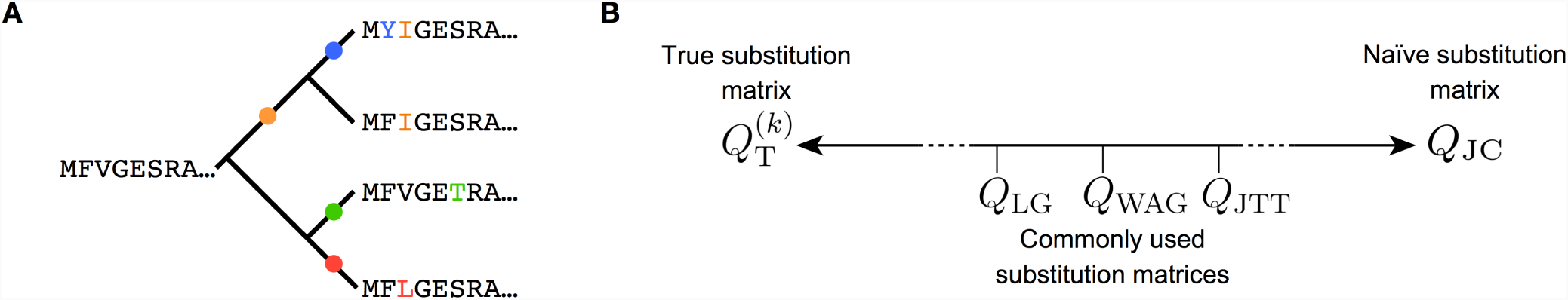
Inference context and framework. (A) We study the process of site-specific rate inference at the amino-acid level in a phylogenetic context. As a protein sequence evolves, substitutions (shown by colored dots and letters) along specific branches introduce site-specific variation. Some sites (e.g., here, site 3) change much faster than others (e.g., sites 1, 4, or 5). The goal of rate inference is to obtain a numeric value that accurately represents the rate at which a site changes. (B) Substitution matrices used for rate inference can have different degrees of realism, indicated along a line from left to right. All the way to the left, we assume there is a true substitution matrix that describes the exact physical process governing the evolution of site *k*. This matrix exists only hypothetically and it can never be known for a natural system. All the way to the right, we have the completely uninformative Jukes-Cantor-like matrix *Q*_JC_, which assumes that all substitution rates are the same for all possible amino-acid substitutions. In between these two extremes reside actual matrices used for rate inference, such as JTT, WAG, or LG.

Here, we specifically study the evolution of protein sequences, and therefore the state space of a Markov chain describing the evolutionary process of an individual site consists of the 20 amino acids. We write the probability of being in the state corresponding to amino acid *i* as *π*_*i*_. State changes from amino acid *i* to *j* occur according to the generator matrix *Q* = *q*_*ij*_, which describes the transition probabilities during an infinitesimally small time interval. For finite times *t*, transition probabilities can be written as *P* (*t*) = *e*^*tQ*^, where *P* (*t*) = *p*_*ij*_ (*t*) is the probability that amino acid *i* is substituted by amino acid *j* during a time interval *t*.

In a conventional data analysis scenario, we will know the sequences from which we want to infer rates but we don’t know the process that guided the evolution of these sequence. Here, to obtain insight into the inference process, we assume that we know the sequence-generating process, and that it can be described by a Markov process as described in the previous paragraph. We will call this model the “true model”, and we denote it with a subscript T, as follows: 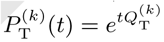. Here, the superscript *k* indicates the site in the sequence described by this model. We assume we have multiple sites, all with their own generating matrix 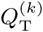. However, *t* is not site dependent. It is shared by all sites. We emphasize that in an actual inference scenario, neither *t* nor *Q*_T_ nor *P*_T_(*t*) would be known.

To measure site-wise rates of evolution from the sequences generated by the true model, we fit a different Markov model to the sequences. We call this model the “measurement model”, and we denote it with a subscript M, as follows: 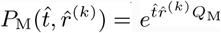. The measurement model is similar but not identical to the true model. First, because we don’t know the elements of the true model matrix *Q*_T_, we cannot guarantee that the measurement matrix *Q*_M_ is identical to the true matrix at any site. In fact, for most of this paper, we assume the same matrix *Q*_M_ is used at all sites. Second, instead of a single scalar parameter *t* describing divergence time/branch length, we have two, 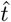 and 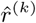. 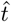 is the estimated branch length, shared among all sites, and 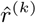 is the estimated evolutionary rate at site *k*. Because the three quantities 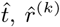 and *Q*_M_ enter the term 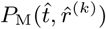 via a product, the choice of *Q*_M_ will generally affect the inferred branch length 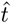 and site-specific rate of evolution 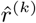. We also note that 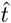 and 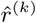 always appear as the product 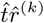 and hence they are only uniquely specified up to a multiplicative constant. We resolve this ambiguity by enforcing the mean of 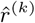 over all sites *k* to be one.

Because our inference model *a priori* makes no specific assumptions about the elements of the measurement matrix 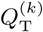, we have a wide range of different choices. We can arrange these choices along a line from most realistic to least realistic when compared to the true matrix. At one end of this line lies the true matrix 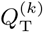 (Figure 1B). At the other end lies the completely uninformative, naïve matrix, which has no knowledge about the substitution rates between amino acids. It assumes that all rates of substitution are equal to each other. Thus, it can be thought of as a Jukes-Cantor-like matrix (36) for amino acids, and we refer to it as *Q*_JC_ (Figure 1B). Commonly used substitution matrices, such as JTT (19), WAG (20), and LG (21) fall somewhere in between these two extremes. They will be more realistic (and hence closer to the true matrix) than *Q*_JC_, but they certainly are not equal to 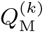, in particular since the true substitution process has to be assumed to be different at each site. Because any matrix *Q*_M_ we may use falls somewhere on this line from completely true to entirely uninformative, we can systematically explore how the choice of measurement matrix influences the estimates of site-wise rates. In particular, we can analytically solve the inference equations at the two extreme ends, and we can also solve the inference equations in the limit of *t →*0. In combination with numerics for cases where an analytic solution is not available, this analysis provides for a comprehensive characterization of the choice of measurement matrix on the inference process.

### Rate measured with an arbitrary measurement matrix for *t* ≪ 1

We assume that the measurement model is fit to the sequence data by maximum likelihood (ML). As is customary, we assume that the measurement model and the true model are time-reversible (29). For analytic tractability, we consider the simplest possible case of two sequences that have diverged for some time *t*. Because site-wise parameters cannot be reliably inferred from an alignment consisting of only two sequences, we employ a mathematical trick and assume each site in the alignment is duplicated *n* times. As part of our derivation, we can show that the choice of *n* is irrelevant, and our results remain valid (on average) in the limit of *n→*1. The detailed mathematical formulation of our theory and detailed derivations are provided in the SI Appendix.

First, we analytically derive site-wise rates measured with an unspecified measurement matrix 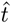 in the limit of *t →* 0. In this limit, we can assume that the estimated time *t* is proportional to the true time *t*, 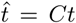 = *Ct* for some constant *C*. Without loss of generality, we set *C* = 1. (We implicitly obtain the correct *C* when we normalize the rates 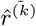. In this scenario, we find (SI Appendix)

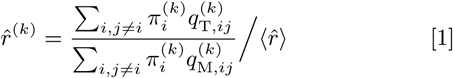

with

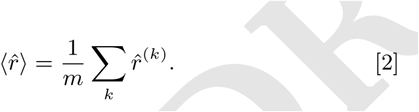

Here, *m* is the total number of sites in the sequence, and 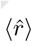 is the mean rate across sites, which we use for normalization. By 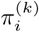 we denote the stationary probabilities of the true model at site *k*. Note that 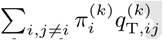 is the steady-state rate of substitutions under the true model at site *k*, and 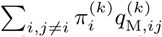 is the rate with which the process would leave the true-model steady state if the transition matrix were switched to the measurement matrix. Thus, the inferred rate 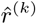 is the ratio of the true steady-state substitution rate to the substitution rate under the measurement matrix, starting from the true equilibrium state.

Eq. (1) provides several key insights. First, if the measurement matrix equals the true matrix, then the relative rate 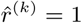. Second, for any measurement matrix for which all the column sums are equal, 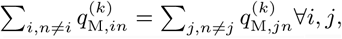, the denominator drops out of the equation and the estimated rate equals the steady-state rate under the true model. Third, when the column sums of the measurement matrix are not equal, the meaning of the inferred 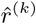 is unclear.

### Rates measured at arbitrary *t*, using the naïve substitution matrix

The two extremes described in Figure 1B, where the measurement matrix is either the true matrix or the naïve matrix, can be further analyzed for any divergence time *t*. If the measurement matrix is the same as the site’s true substitution matrix, then the true model and the fitted model become the same. In this case, the inference model recovers true time 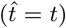 and the estimated rate becomes 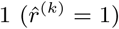. So, for any site, if inference is performed using the true substitution matrix, then inference of a relative rate is meaningless, since that rate will always be measured as 1.

Next we consider the case where the site-wise rate is measured using the naïve substitution matrix, which we also refer to as the Jukes-Cantor-like matrix (JC). In this matrix, the substitution rate from any amino acid to any other amino acid is 1/19. After solving the ML equations, we find (SI Appendix)

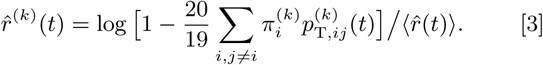

We note that this equation depends on *t*; the estimated rate is not constant. In the limit of *t →* 0, the estimated rate becomes

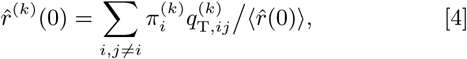

consistent with Eq. (1).

For completeness, we also report the limit of Eq. (3) for *t → ∞*, which is

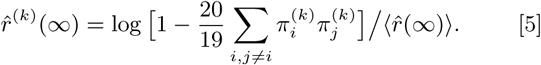

To test our theory numerically, we simulate sequences under a given true model and then infer rates via the standard maximum likelihood approach. To generate realistic simulated sequences, we use a mutation–selection (MutSel) model (38,39) which we parameterize from predicted stability effects of mutations in a protein structure (37). We find that when we use *Q*_JC_ in the maximum likelihood inference, as assumed in the derivation of equations 3–5, the predicted rate estimates agree perfectly with the average inferred rate estimates, at all times *t* (Figure 2).

**Fig. 2.**
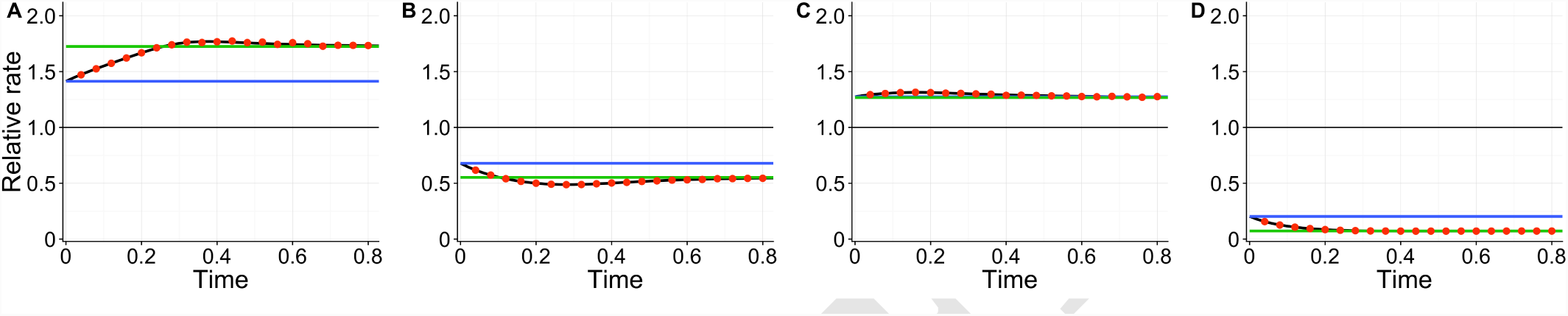
Comparison of the inferred rates and the analytically derived rates, in a model of two sequences diverged for time *t* and with *n* = 100, 000 site duplicates. Rates are inferred with the Jukes-Cantor-like matrix and normalized to their mean across sites. The black line represents the analytically derived rate for arbitrary time *t* (equation 3). The blue line represents the analytically derived rate when time *t* is small (equation 4). The green line represent the analytically derived rate when time *t* is large (equation 5). The red dots represent the mean inferred rate at each time point across 30 simulations. The error bars represent the standard error. For all points, the bars are smaller than the symbol size. The horizontal line at 1 represents the average rate in the sequence. (A-D) Rate over time for sites 1, 2, 4, and 5, respectively, in egg white lysozyme (PDB ID: 132L)(37). Convergence to the predicted mean rate with increasing number of site duplicates *n* is shown in Fig. S1.

### Measurement using a matrix that is proportional to the true matrix

As equation 3 shows, in the general case of measurement using the JC-like matrix the estimated rates change with divergence time *t*. Only in the limit of small *t* (equation 4) do we recover the result that the estimated rate corresponds to the mean substitution rate in the true model. To investigate the cause of this time dependency, we derive the estimated rate under the assumption that the measurement matrix is proportional to the true matrix. Specifically, we assume that the true matrix 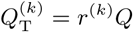, where *Q* is any arbitrary substitution matrix, and we then set 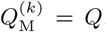. In this case, we can show that the fitted model recovers true time 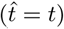 and the true rate parameter modulating 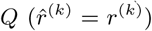 (SI Appendix).

To numerically validate this calculation, we simulate evolution using a true model with 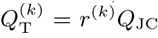 and the *r*^(*k*)^ at each site drawn from a gamma distribution (see Methods). After performing rate inference with 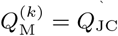, we recover the original *r*^(*k*)^ and the estimates do not depend on *t* as predicted (Figure S2).

These results suggest that the time dependence of the inferred rate seen in equation 3 and Fig. 2 is caused by the mismatch between the true model and the JC inference model. Even though the rate should be a separate parameter independent of *t*, when we use a JC inference model the inferred rate depends somewhat on divergence time. However, this artifact disappears when the measurement matrix is proportional to the true matrix, in which case ML can recover the true model parameters. Of course, in any real-world application it is never possible to have the measurement matrix be exactly proportional to the true model matrix, and hence moderate time dependence of the inferred rate parameter has to be expected in all realistic rate-inference applications.

### Rate measured with commonly used matrices

We have been able to analytically solve the maximum likelihood equations for the JC-like matrix, but in practice that matrix is rarely used for rate inference. Instead, commonly used matrices include matrices such as JTT, WAG, and LG, as shown in Figure 1. Moreover, we have solved the ML equations for a pair of diverged sequences and with site duplicates, but in a realistic application we would perform inference on individual sites in a multiple sequence alignment and corresponding phylogenetic tree. To assess the extent to which matrix choice affects rate inference, we use the JTT, WAG, and LG matrices in addition to the JC-like matrix for inference on simulated multiple sequence alignments without site duplicates. For reference, we compare the inferred rates to the predicted rates at small *t* (equation 4), which we hypothesize to generalize to inference on multiple sequences. We find that the rates inferred with JTT, WAG, and LG behave very similar to each other and to the JC matrix for all branch lengths (Figures 3 and S3). In addition, and as expected from the results in the preceding section, the inferred rates are also similar to the analytically derived rates. In aggregate, these results are consistent with a recent study on the impact of matrix choice on rate inference, which has found that inferred rates are largely independent of the chosen matrix (16).

Any general time-reversible model can be decomposed into a symmetric matrix called the *exchangeability matrix* and a vector of equilibrium frequencies (29). Strictly speaking, the various matrices JTT, WAG, and LG are exchangeability matrices that need to be combined with equilibrium frequencies to specify the full model matrix. If we were to use any of these matrices as is, we would implicitly assume uniform amino-acid frequencies. This is not normally done, however. Instead, the observed amino-acid frequencies in the multiple sequence alignment are commonly used as equilibrium frequencies, and we follow this practice here for JTT, WAG, and LG (see Methods). By contrast, our theoretical derivation with the JC-like matrix uses the matrix as is and thus assumes uniform frequencies in the measurement model. To assess the impact of this choice, we also perform inference with the JC-like matrix and observed frequencies. We find that the choice of assumed equilibrium frequencies in the measurement model has virtually no effect on the inferred rates (Figure S4).

### Codon rates measured with a naïve amino acid substitution model

Gene sequences in natural organisms do not evolve according to an amino acid matrix *Q*. Protein sequences are encoded in DNA, and the DNA is the substrate that mutates and evolves. Yet inference models are frequently formulated as amino-acid models. To investigate the effect of this model mismatch, we derive expected inferred rates when the true model is a codon model and the inference model is an aminoacid model using the JC-like matrix (SI Appendix). The derivation follows the same logic as before, and for small *t* we find

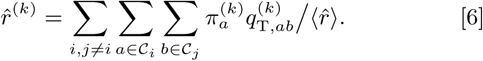

Here, 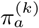 is the true codon equilibrium frequency for a codon *a* and 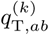 is the true substitution rate between codons *a* and *b*. 𝒞_*i*_ is the set of codons that encode amino acid *i*, and the first sum (over *i* and *j*) runs over all possible amino-acid pairs. As before, our analytic predictions of inferred mean rate agree well with simulations (Figure S5).

Equation 6 demonstrates that the inferred rate reproduces (on average) the true mean substitution rate between all nonsynonymous codons. Thus, it is meaningful to perform an amino-acid based inference on codon-level data. However, because this inference only considers changes between amino acids, we cannot infer rates between synonymous codons.

## Discussion

The conventional inference approach in molecular evolution consists of fitting a series of models by maximum likelihood or Bayesian methods, identifying the best fitting model via likelihood ratio test or some information criterion, and then taking the inferred parameters of the best fitting model as the best estimate for the parameter(s) of interest. However, an increasing body of work shows that this process can go wrong when the model is misspecified (39–42). In particular, the estimates for parameters of biological interest can be much closer to the true value in poorly fitting models (as evaluated by an information criterion) than in the best fitting models (39). A recent paper has proposed the term *phenomenological load* to describe this effect (41). On the flip side, models with very different information scores can also produce substantively similar parameter estimates (16). We believe that the approach we have described here, which relies neither on good model fit nor on proper model specification but instead treats the model fitting as a measurement process, can provide a solution to the problems of model misspecification, phenomenological load, and model selection. If we know what kind of a measurement a model performs on a given dataset, we immediately obtain an interpretable result, from one fit of one single model. For example, in the case described here, measurement with the Jukes-Cantor-like matrix returns rates that correspond to the mean substitution rate in the true model. Of course, as our calculations have shown, this statement is strictly true only in the limit of low sequence divergence. However, the systematic deviations for higher sequence divergence tend to be moderate, in particular when compared to the measurement noise inherent in making singe-site estimates, and hence we believe that making rate measurements with the Jukes-Cantor-like matrix is a principled approach that is as good as or better than the alternatives at any sequence divergence level.

In phylgenetic inference, it remains an open question which empirical matrix is the most appropriate, though LG tends to produce better likelihood scores than do JTT or WAG (21). We have seen here, however, that in the context of rate inference, the rates inferred by LG, JTT, WAG, or JC are nearly identical. A similar observation was recently published in an empirical survey of different approaches to rate inference (43). While JC is uninformative whereas LG, JTT, and WAG are derived from natural protein sequences, we believe that all these matrices are sufficiently similar to each other that they are nearly identical for inference purposes. The explanation for the matrices’ similarities lies in how they were derived and what they represent. JC assumes that any amino acid is permissible at any site, i.e., it assumes a completely flat fitness landscape. JTT, WAG, and LG allow almost as wide of a range of amino acids at each site (44), because they have been derived by pooling data from many sites in many proteins, thus averaging over the fitness differences between different amino acids at different sites and not actually representing any specific site in a protein (45). While one could conclude from these observations that the choice of the inference matrix is irrelevant (43), we believe that JC has a clear theoretical advantage over the other matrices: Equation 4, which states that in the limit of small divergence time *t* the inferred rate converges to the mean substitution rate in the true model, is generally not true for the other matrices, because of equation 1. The denominator of 1 cancels for the JC-like matrix but it does not do so for the empirically derived matrices.

The model we have discussed here for measuring site-specific rates has a separate rate parameter at each site, and thus is a fixed-effects model. While some existing inference tools similarly implement fixed-effects inference (43), others fit a rate distribution to all sites at once, in a random-effects framework (14, 15). A priori, our theory does not apply to random-effects inference. However, empirically it is found, across a variety of different modeling scenarios and frameworks, that fixed-effects approaches and random-effects approaches yield rates estimates that are highly correlated (43, 46, 47). Therefore, we expect that our theoretical predictions will be approximately correct for inference approaches based on random-effects frameworks as well.

In our theory, evolutionary distances between amino acid sequences are measured by the site-wise rate 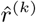 and time *t*. In all derivations, these two quantities appear as the product 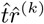. This product is invariant to a rescaling of 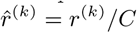 and 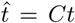, and rate, therefore, is not unique. To obtain unique rates, we have to impose a normalization condition. We have here normalized site-wise rates by the average rate among all sites of the sequence, 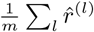. This normalization allows for simple interpretation of the rate as the relative increase or decrease compared to the average rate of evolution in the sequence. Alternatively, we could normalize rates to the median, which may be advantageous when rate distributions are overdispersed (43).

We note several limitations to our findings. First, our calculations assume that sites evolve independently of each other. Yet epistatic interactions among protein sites are widespread, in particular among sites that are in direct contact in the 3D structure (48–51). Strictly speaking, our results do not hold for interacting sites. However, we note that in our derivation, interacting sites would only enter the true model, not the measurement model. Thus, to the extent that the true evolutionary dynamic (with interactions) generates a stochastic process at individual sites that looks Markovian, our results should carry over. Future work will have to test this hypothesis more systematically. Second, in our simulations, we used uniform mutation rates, so that the site-specific variation in evolutionary rates was fully determined by fitness differences among amino acids or codons. For more complex mutation schemes, the inferred rates would be influenced by both the mutation scheme and the fitness differences, and thus, with the measurement process we describe here it is not possible to disentangle mutation and selection. However, we emphasize that similar problems arise in other inference approaches (41) as well, and that the rate equations we have derived are correct regardless of the chosen mutation scheme. Finally, our entire analysis is true only on average and/or in the limit of infinitely large samples. We have ignored what (if any) effect the choice of the measurement matrix has on the variation in the estimate at finite sample sizes. Numerically, it appears that some matrix choices result in less variable estimates than others (Figure 3). Future work will have to address whether different measurement matrices generate different amounts of variability in rate estimates.

**Fig. 3.**
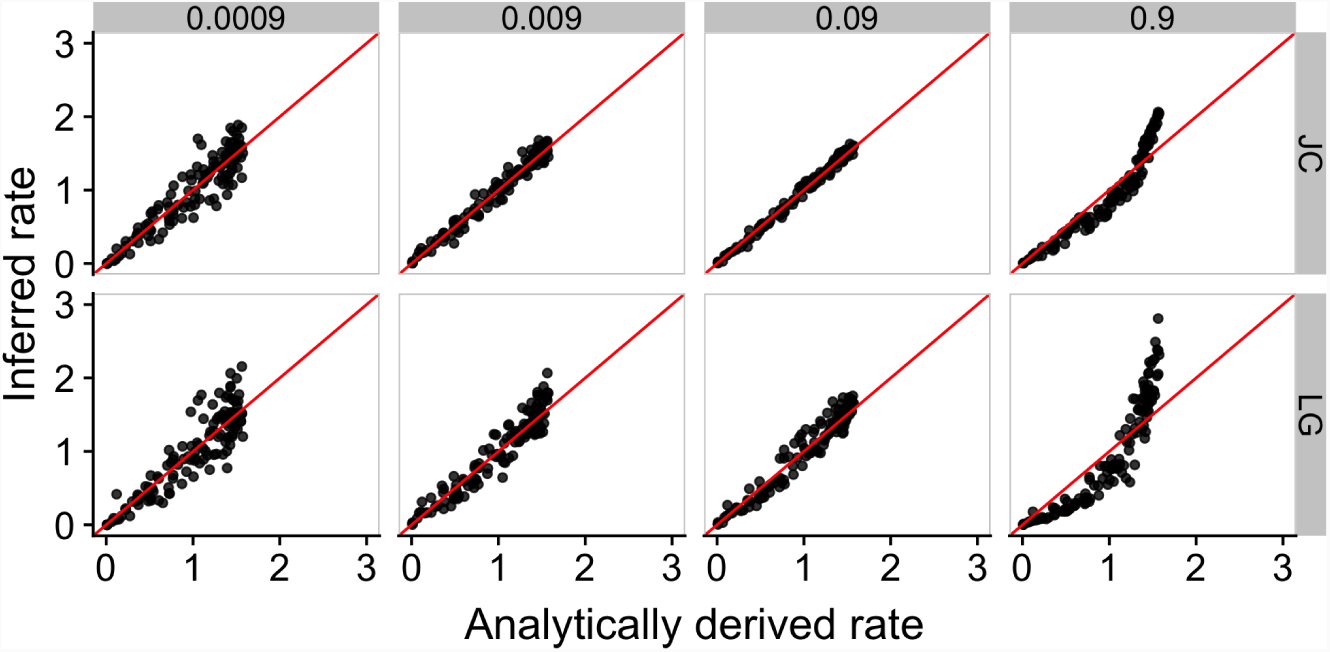
Relationship between analytically derived rates and rates inferred with either the Jukes-Cantor-like (JC) matrix (top row) or the LG matrix (bottom row). Sequence alignments were simulated for binary trees with 512 taxa and 124 sites, parameterized using data from egg white lysozyme (see Methods). No site duplicates were used in these simulations. The inference with LG matrix assumed that each amino acid’s equilibrium frequency is equal to the frequency of that amino acid in the entire alignment. The inference with JC matrix assumed that each amino acid’s equilibrium frequency is 1/20. The inferred rates plotted above are mean inferred rates over 50 simulations for each time point and site. The analytically derived rate was calculated with equation 4. The numbers on top of the plot panels indicate the time *t* used for each simulation. The labels on the right indicate the substitution matrix used for inference. Each point represents one site, and the diagonal line represents *x* = *y*.

## Materials and Methods

For detailed mathematical derivations, see SI Appendix.

### Parameterizing a MutSel model using stability effects of mutations in a protein

To parameterize a MutSel model for simulation, we need to define site-specific scaled selection coefficients 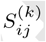 for all pairs of amino acids *i* and *j*. To arrive at a somewhat realistic parameterization, we used a recently proposed framework to link protein stability to protein evolution (37). Under this framework, we can write (SI Appendix)

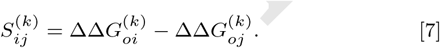

Here, 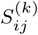 is the scaled difference in fitness between amino acid *i* and amino acid *j* at site *k* and 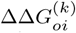 is the change in stability of the protein structure when a reference amino acid (which can be arbitrarily chosen) is substituted with amino acid *i* at site *k*. We used 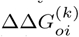 data for egg white lysozyme (PDB ID: 132L) from Ref. (37), and we calculated scaled selection coefficients for all sites for which ΔΔ*G* values were available (124 sites in total). From the selection coefficients, we then calculated a substitution matrix *Q* following standard MutSel theory (38) (see SI Appendix for details).

### Generation of simulated sequence alignments

In all cases, we first generated binary trees with chosen branch lengths and numbers of taxa, using the R package ape (52). We then evolved sequences along each tree using the python library pyvolve (53), and each site was evolved according to a site-specific MutSel model. The MutSel models were parameterized according to equation 7 as described above, except in the case where 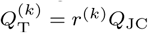. In this latter case, we did not have to choose scaled ^T^selection coefficients but instead values of the true rate *r*^(*k*)^ at each site. We assumed that the true rates are gamma distributed, as is commonly observed in empirical work, and we randomly drew true rates from a gamma distribution with shape parameter *α* = 0.312 and rate parameter *β* = 1.027. These are parameter values that have been estimated for HIV-1 integrase (Table 2 in (54)).

#### Generating amino-acid alignments to verify rate derivations

Our simulations directly mimicked the assumptions made in the analytical derivations. We simulated two sequences that have evolved from a common ancestor and that have diverged over time. We took the first ten sites from the egg white lysozyme and duplicated each site *n* times. The site duplicates were simulated to all evolve independently under identically parameterized models.

In a first set of simulations, we set *n* = 100, 000 and varied branch lengths across 25 different values. Each tree in our simulations had 2 branches of equal lengths, and the length of the branches ranged from 0.02 to 0.5 in the increments of 0.02 (this is equivalent to a time range from 0.04 to 1 in the increments of 0.04). We simulated 30 replicates per tree, for a total of 750 alignments.

In a second set of simulations, we set branch length along the two branches to 0.24 each (*t* = 0.48) and varied *n* from 10 to 100,000 in multiples of 10. We simulated 50 replicates per number of site duplicates, for a total of 250 alignments.

#### Generating codon alignments to verify rate derivations

For the case when the true model is a codon model, we simulated two codon sequences according to a codon MutSel model. Each site was simulated according to a codon substitution matrix, which was derived and parameterized similarly to the amino-acid case. We introduced codon scaled selection coefficients 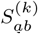 analogously to equation 7. We set codon ΔΔ*G* values equal to the ΔΔ*G* of the amino acid the codons translate to. The 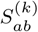 between two synonymous codons *a* and *b*, therefore, was 0. The 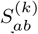 between two non-synonymous codons *a* and *b* was set to equal the difference in the ΔΔ*G* values between the two codons. We set the mutation rate between any two codons to 1. As in the amino-acid case, we simulated 10 sites of egg white lysozyme with *n* = 100, 000 duplicates each. The sequences were evolved along 25 different trees. Each tree had 2 branches of equal lengths, and the length of the branches ranged from 0.02 to 0.5 in the increments of 0.02 (this is equivalent to a time range from 0.04 to 1 in the increments of 0.04) with one tree per branch length value. We simulated 30 replicates per tree, for a total of 750 alignments.

#### Generating alignments to compare rates inferred with JTT, WAG, LG, and JC matrices

We generated 4 binary trees with four different branch lengths (0.00005, 0.0005, 0.005, and 0.05) and with 512 taxa each. We set the number of taxa to 512 because higher numbers of taxa reduces the variation in inferred site-wise rates (55). We simulated amino acid sequences with 124 sites parameterized according to the ΔΔ*G* values for egg white lysozyme, as described above. No site duplicates were simulated, so that each site appeared in each sequence exactly once. We simulated 50 replicates per tree, for a total of 200 simulated alignments.

### Rate inference

Similarly to our simulation approach, our inference approach was intended to closely follow the assumptions made in the analytical derivations of site-wise rate. We inferred rates and time (branch length) from simulated sequences via HyPhy (11), using code similar to the recently published LEISR model (43). For all simulations, before inferring site-wise rates, we inferred time 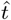 by fitting a model with 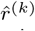 set to 1 jointly to all sites in the alignment. Subsequently, we inferred site specific rates by fitting a separate model at each site by maximum likelihood while fixing 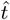 to the previously inferred global value.

To confirm rate derivations for equations 3–5, we performed inference using the Jukes-Cantor-like (JC) matrix (every off-diagonal element is 1/19 and every element on the diagonal is 1) with uniform amino-acid equilibrium frequencies (the frequency of each amino acid is equal to 1/20). We inferred one rate per site, with the *n* duplicates counting as one joint site, for a total of 10 inferences per 10 simulated sites. The rates were normalized by the average inferred rate in the sequence.

To assess the effect of other measurement matrices, we performed similar inferences on the simulated alignments. For each simulated alignment, we inferred site-wise rates five times, once each for the matrices JTT, WAG, and LG, and twice for the JC-like matrix, using different assumptions of equilibrium amino acid frequencies. For JTT, WAG, and LG, the equilibrium frequencies were assumed to be equal to the observed amino-acid frequencies. For JC, we inferred rates once under the same assumption about equilibrium frequences, and once under the assumption that the equilibrium frequency for each amino acid is 1/20, the assumption made in the theoretical derivations. Both assumptions yield almost identical rates (Figure S4). We inferred one rate per physical site (no duplicates were simulated). For each alignment, the final rates were normalized by their mean.

When the true model was assumed to be a codon MutSel model, we translated codon sequences to amino acid sequences prior to inference. To match branch lengths between codon and amino-acid alignments, we multiplied codon times by 0.77, the expected number of amino-acid substitutions per nucleotide substitution according to the genetic code. We inferred site-wise rates with the JC-like matrix and uniform amino-acid frequencies. For each site, one joint rate was inferred from the site’s duplicates. For each alignment, the final rates were normalized by their mean.

### Data availability

Code and processed data can be found at https://github.com/dariyasydykova/rates_measurement.

## ACKNOWLEDGEMENTS

This work was funded by National Science Foundation Cooperative Agreement no. DBI-0939454 (BEACON Center) and National Institutes of Health grant R01 GM088344. We thank Julian Echave and Stephanie Spielman for stimulating discussions on this topic.

